## Supplementary Information for "ssDNA recombineering boosts *in vivo* evolution of nanobodies displayed on bacterial surfaces"

**Supplementary Table S1.** Strains used in this work

| Strain *Escherichia coli* | Description | Reference |
|---|---|---|
| DH5α | *supE44, ΔlacU169 (φ80 lacZΔM15), hsdR17, recA, endA1, gyrA96,* $thi^{-1}$*, relA1* | [1] |
| BL21(DE3) | *hsdS, gal* (*λ*$cI^{ts}$*857, ind1, Sam7, nin5, lacUV5-T7* gene 1) | [2] |
| HB2151 | *nalr thi-1 ara ∆(lac-proAB) [F' proAB+ lacIq lacZ(M15]* | [3] |
| WK6 | F' lacIq ∆(lacZ)M15 proA+B+ ∆(lacproAB) galE rpsL | [4] |
| EcM1luxSATir | MG1655∆fimA-H ∆flu::$P_{N25}$-SAtir ∆matB::$P_2$-luxCDABE | [5] |
| EcM1luxSATir<br><br>(pORTMAGE-3) | Bacterial strain that expresses constitutively the Nb TD4 and enables its display in the cell surface. This strain was used as parental for the generation of libraries. The plasmid pORTMAGE-3 is to enhance the efficiency of recombineering events. | This work |
| BL21 DE3 (pET28a_TirM EPEC) | Used for the overproduction of the antigen EPEC TirM. | |
| BL21 DE3 (pET28a_TirM EHEC) | Used for the overproduction of the antigen EHEC TirM. | [4, 6] |
| EcM1luxNbH107Y | Strain isolated from the library. Bears the cassette in the genome for the expression of the Nb with the replacement H107Y. | This work |
| EcM1luxNbT108R | Strain isolated from the library. Bears the cassette in the genome for the expression of the Nb with the replacement T108R. | This work |
| EcM1luxNbD116G | Strain isolated from the library. Bears the cassette in the genome for the expression of the Nb with the replacement D116G. | This work |
| HB2151(pVdL9.3_Vamy) | Strain that bears the plasmid for the overexpression of the Nb Vamy, which we used as the unspecific control antibody. | [7] |
| HB2151(pVdL9.3_TD4) | Strain that bears the plasmid for the overexpression of the Nb TD4. | |
| HB2151(pEHLYA_H107Y + pVdL9.3) | Strain carrying the plasmids for the expression and secretion of the Nb H107Y. | This work |
| HB2151(pEHLYA_T108R + pVdL9.3) | Strain carrying the plasmids for the expression and secretion of the Nb T108R. | This work |

| | | |
|---|---|---|
| HB2151(pEHLYA_D116G + pVdL9.3) | Strain carrying the plasmids for the expression and secretion of the Nb D116G. | This work |
| (pCANTAB6_VhhTD4) | Produces the plasmid pCANTAB6_VhhTD4 used afterwards as template to obtain, by site directed mutagenesis, the different Nb gene variants which were cloned in pVdL9.3 plasmid and expressed in HB2151. | [4] |
| WK6 (pCANTAB6_H107Y) | Used to obtain the plasmid pCANTAB6_H107Y that was digested SfiI/NotI to clone the gene of the Nb H107Y in pVdL9.3 plasmid. | This work |
| WK6 (pCANTAB6_T108R) | Used to obtain the plasmid pCANTAB6_T108R that was digested SfiI/NotI to clone the gene of the Nb T108R in pVdL9.3 plasmid. | This work |
| WK6 (pCANTAB6_D116G) | Used to obtain the plasmid pCANTAB6_D116G that was digested SfiI/NotI to clone the gene of the Nb D116G in pVdL9.3 plasmid. | This work |

**Supplementary Table S2.** Plasmids used in this work

| Name | Description | Reference |
|---|---|---|
| pORTMAGE3 | Plasmid for the transient expression (controlled by temperature sensitive cI857 repressor and pL promoter) of λ Red Exo, Beta, Gam, and the MutL E32K protein, which confers a dominant mutator phenotype, to enhance the establishment of nucleotide replacements in the genome during DIvERGE. Confers resistance to kanamycin. | [8] |
| pET28a-TirMEHEC | For the overproduction of the antigen EHEC TirM and subsequent purification. Confers resistance to kanamycin. | [4] |
| pET28a-TirMEPEC | For the overproduction of the antigen EPEC TirM and subsequent purification. Confers resistance to kanamycin. | [4] |
| pCANTAB6_VHHTD4 | Bears the gene for VHHTD4. This plasmid was used as template for the transplantation of the genes for new Nb variants by codon replacement. Bears a pUC-ori and confers resistance to ampicillin. | [4] |
| pEHLYA5_Vamy | Used for the purification of the Nb Vamy, specific for alpha amylase. Confers resistance to ampicillin. | [9] |
| pCANTAB6_NbH107Y | Used as source of the gene encoding the NbH107Y by digestion with SfiI and NotI. Confers resistance to ampicillin. | This work |
| pCANTAB6_NbT108R | Used as source of the gene encoding the NbT108R by digestion with SfiI and NotI. Confers resistance to ampicillin. | This work |
| pCANTAB6_NbD116G | Used as source of the gene encoding the NbD116G by digestion with SfiI and NotI. Confers resistance to ampicillin. | This work |
| pVdL9.3 | Encodes HlyB and HlyD components of the haemolysin secretion system, for controlled expression under the Plac promoter. pSC101-ori. Confers resistance to chloramphenicol. | [7] |
| pVdL9.3_Vamy | Used for the purification of the Nb Vamy, specific for alpha amylase, from E. coli culture supernatants using the haemolysin secretion system. Confers resistance to chloramphenicol. | [7] |
| pVdL9.3_TD4 | Used for the purification of the Nb TD4, specific for the antigen TirM EHEC, from E. coli culture supernatants using the haemolysin secretion system. Confers resistance to chloramphenicol. | [7] |
| pEHLYA_NbH107Y | Used in combination with the plasmid pVdL9.3 for the purification of the NbH107Y from E. coli culture supernatants using the haemolysin secretion system. | This work |
| pEHLYA_NbT108R | Used in combination with the plasmid pVdL9.3 for the purification of the NbT108R from E. coli culture supernatants using the haemolysin secretion system. | This work |

| | | |
|---|---|---|
| pEHLYA_NbD116G | Used in combination with the plasmid pVdL9.3 for the purification of the NbD116G from E. coli culture supernatants using the haemolysin secretion system. | This work |

**Supplementary Table S3.** Oligonucleotides used in this work

| Oligo ID | Name | Sequence / description | Tm ºC |
|---|---|---|---|
| 1 | TD4_T108R_F | CATGGGACCGCGCCATATTGGCACAgGCCCATCCCTACTCTCTCCGAAG | 87 |
| 2 | TD4_T108R_R | CTTCGGAGAGAGTAGGGATGGGCcTGTGCCAATATGGCGCGGTCCCATG | 87 |
| 3 | H107Y_Fwd | GACCGCGCCATATTGGtACACGCCCATCCCTACT | 79 |
| 4 | H107Y_Rev | AGTAGGGATGGGCGTGTaCCAATATGGCGCGGTC | 79 |
| 5 | D116G_Fwd | CCCTACTCTCTCCGAAGgTAAGTATTTCTACTGGG | 75 |
| 6 | D116G_Rev | CCCAGTAGAAATACTTAcCTTCGGAGAGAGTAGGG | 75 |
| 7 | pCANTAB6_F | CCAGTACACTCCTGTATCATCAAAAGCC | 68 |
| 8 | pCANTAB6_R | CTCTTCGCTATTACGCCAGC | 60 |
| 9 | CDRI_Fwd | CTCAGGTGCAGCTGGTGGACG | 67 |
| 10 | CDRII_Midd_R | CAGGCTGTTCATCTGCAGATATATCGTG | 68 |
| 11 | CDRIII_Rev | CAGCTGCATCCTCTTCTGAGATGAG | 67 |
| 12 | CheckSAtir_F | CGCCTATGACCGTAATGGCAATAGCTCTAACAATGTACAGC | 78 |
| 13 | CheckSAtir_R | TCAGAAGGTCACATTCAGTGTGGCCTGACCGTTATACC | 78 |
| 14 | tirDSF | CTCGGTGCAGGCTGGA | 56 |
| 15 | tirDSR | CCGCTGAGGAGACGGTGACCTG | 69 |
| 16* | tirRM1 | GGGTCTCTAACACTCTCCTGTGTAGCCTCTGGAGCCGCCTACAGTACGAACTTGTTGGGCTGGTTCCGCCAGGCTCCAGGGAAGGAGCGC | - |
| 17* | tirRM2 | GGGGTCGCATCTATTTATCGTGGTAATAGTGCCACGAACTATGCCGACTCCGTGAAGGGCCGATTCACCATCTCCCAAGACAAGACCAAA | - |
| 18* | tirRM3 | CATGTACTACTGTGCACATGGGACCGCGCCATATTGGCACACGCCCATCCCTACTCTCTCCGAAGATAAGTATTTCTACTGGGGCCAGGG | - |

*underlined sequence regions were synthesized with soft-randomized monomer mixture as follows: A = 98.5%A+0.5%C+0.5%T +0.5%G; C = 98.5%C+0.5%A+0.5%T +0.5%G; G = 98.5%G+0.5%C+0.5%T +0.5%A and T = 98.5%T+0.5%C+0.5%A +0.5%G.

**Supplementary Table S4.** Amino acid sequence of the different nanobodies

| | |
|---|---|
| Nb TD4 | MAQVQLVDAGGGSVQAGGSLTLSCVASGAAYSTNLLGWFRQAPGKEREGVASIYRGNS ATNYADSVKGRFTISQDKTKYTIYLQMNSLKPEDSAMYYCAHGTAPYWHTPIPTLSEDKYF YWGQGTQVTVSSAA |
| Nb H107Y | MAQVQLVDAGGGSVQAGGSLTLSCVASGAAYSTNLLGWFRQAPGKEREGVASIYRGNS ATNYADSVKGRFTISQDKTKYTIYLQMNSLKPEDSAMYYCAHGTAPYWYTPIPTLSEDKYF YWGQGTQVTVSSAA |
| Nb T108R | MAQVQLVDAGGGSVQAGGSLTLSCVASGAAYSTNLLGWFRQAPGKEREGVASIYRGNS ATNYADSVKGRFTISQDKTKYTIYLQMNSLKPEDSAMYYCAHGTAPYWHRPIPTLSEDKYF YWGQGTQVTVSSAA |
| Nb D116G | MAQVQLVDAGGGSVQAGGSLTLSCVASGAAYSTNLLGWFRQAPGKEREGVASIYRGNS ATNYADSVKGRFTISQDKTKYTIYLQMNSLKPEDSAMYYCAHGTAPYWHTPIPTLSEGKYF YWGQGTQVTVSSAA |

**Supplementary Figure S1.** Protein verification by SDS-PAGE

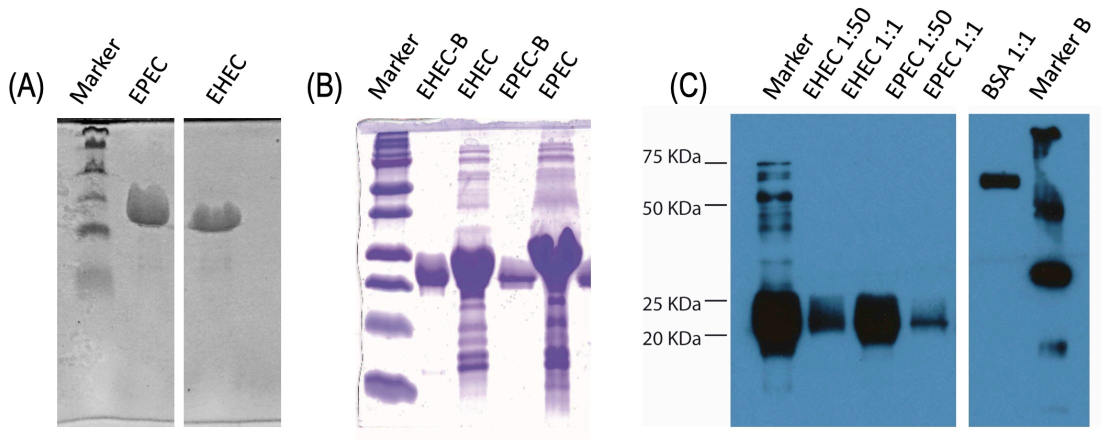

The antigens EPEC and EHEC were checked during the different preparation steps. (A) SDS-PAGE after purification by Immobilized Metal Affinity Chromatography (IMAC) and dialysis. Marker band sizes are 7.1, 20.6, 28.9, 34.8, 49.1, 80, 124, and 209 kDa. (B) SDS-PAGE with samples of the antigens biotinylated, EHEC-B and EPEC-B, and without biotinylation, EHEC and EPEC. (C) Western blot showing different sample dilutions of the biotinylated antigens.

**Supplementary Figure S2.** Effects of the enrichment process

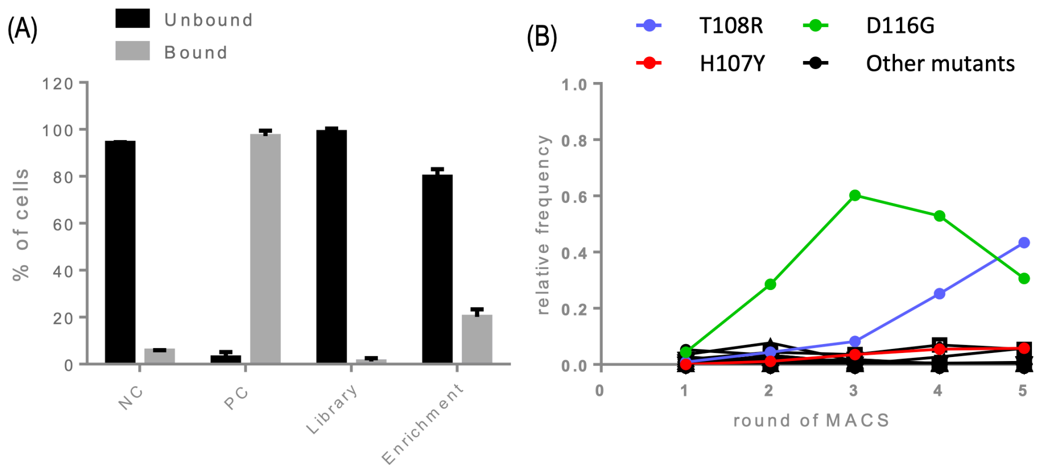

(A) NC corresponds to a strain that expresses and displays a nanobody that is specific for GFP recognition and it is used as unspecific control for this experiment. PC (positive control) corresponds to the parental nanobody TD4 and it is used as control of a positive interaction. Almost all of the cells that display the parental nanobody can bind specifically to the former antigen (PC), but nevertheless, the number of cells from the library that are able to bind the new antigen is extremely low (Library). In contrast, after the enrichment, the bias towards the ability to bind the new antigen becomes evident as shown by the increase in cells that bind to EPEC TirM after the enrichment (Enrichment). (B) Based on mutant counts found by sequencing with PacBio technology. The frequency of mutant clones after each round of library enrichment is represented (number of times a sequence appeared/number of times all the mutant sequences appeared). Compared to the rest of the mutants in the library, the abundance of clones producing any of the nanobody T108R and D116G is specially increased during the enrichment. The trend is represented in colors for the three clones that were isolated for deeper characterization in this work and in black for the rest of mutants.

**Supplementary Figure S3.** Verification of the Nb variants by SDS-PAGE

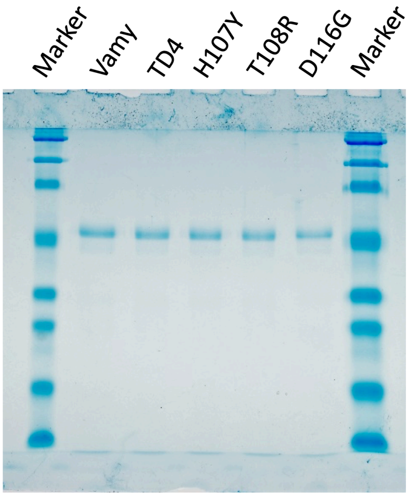

Nb variants after they were purified and concentrated were checked before they were used in ELISA. From each Nb sample 6 µg were loaded. A volume of 5 µL and 10 µL of marker were loaded on the left and right well respectively. Marker band sizes are 7.1, 20.6, 28.9, 34.8, 49.1, 80, 124, and 209 kDa.
